# Gene drive escape from resistance depends on mechanism and ecology

**DOI:** 10.1101/2021.08.30.458221

**Authors:** Forest Cook, James J. Bull, Richard Gomulkiewicz

## Abstract

Gene drives can potentially be used to suppress pest populations, and the advent of CRISPR technology has made it feasible to engineer them in many species, especially insects. What remains largely unknown for implementations is whether anti-drive resistance will evolve to block the population suppression. An especially serious threat to some kinds of drive is mutations in the CRISPR cleavage sequence that block the action of CRISPR, but designs have been proposed to avoid this type of resistance. Various types of resistance at loci away from the cleavage site remain a possibility, which is the focus here. It is known that modest-effect suppression drives can essentially ‘outrun’ unlinked resistance even when that resistance is present from the start. We demonstrate here how the risk of evolving (unlinked) resistance can be further reduced without compromising overall suppression by introducing multiple suppression drives or by designing drives with specific ecological effects. However, we show that even modest-effect suppression drives remain vulnerable to the evolution of extreme levels of inbreeding, which halt the spread of the drive without actually interfering with its mechanism. The landscape of resistance evolution against suppression drives is therefore complex, but avenues exist for enhancing gene drive success.

## Introduction

Synthetic gene drives are poised to have profound effects on the way humans control pests and pathogens. Not only can they be used to quickly transform wild populations with genes of social value, even if detrimental to the species, but they can be used to suppress and even extinguish populations (Hamilton 1967; Lyttle 1977). The extinction potential has been known for over half a century, with the remaining hurdle to implementation having been engineering (Burt 2003). CRISPR has solved that hurdle, and there are now countless examples in which gene drives have been genetically engineered, tested, and shown to work in laboratory populations of selected insects (Bier 2021).

What remains is to understand whether suppression drives can escape resistance evolution. The fitness consequences of a suppression drive create an advantage for genes that block the drive, and if those blocks evolve, the suppression drive can be rendered ineffective (Lyttle 1979, 1981). If a suppression drive is to have a lasting effect, especially cause extinction, it either needs to be designed so that resistance is mutationally unlikely or not tolerated (Kyrou et al. 2018; Champer et al. 2018) or that it evolves to fixation ahead of resistance – which requires that the resistance be genetically unlinked from the drive (Gomulkiewicz et al. 2021).

### Realms of resistance

In understanding gene drive evasion of resistance evolution, it is necessary to contemplate the different types of resistance that may occur, because the type of resistance affects the evolutionary dynamics. We can identify three distinct types of resistance (see also Bull et al. 2019)):

‘M’ **Mechanistic.** The drive mechanism is blocked in the individual, impairing the drive’s selfish advantage. With CRISPR, this could take the form of a mutation at the endonuclease cleavage site that blocks cutting or a protein that interferes with CRISPR function (Taxiarchi et al. 2021; Jia and Patel 2021). Some of these would necessarily be allelic to the drive, others could be unlinked.
‘C’ **Compensatory.** The drive evolves normally but the fitness consequence of the drive is abrogated by evolution at another site, such as compensatory evolution elsewhere in the genome (Lyttle 1981; Burt 2003).
‘P’ **Population structure.** The population develops a structure that enables family or group selection to keep the drive from spreading/fixing (Bull 2016; Bull et al. 2019). The population structure need not evolve *per se* but may instead arise dynamically as the population size declines (Bull 2016; Bull et al. 2019; North et al. 2019; Champer et al. 2021).

The practicality of avoiding these alternative resistances will depend on the application as well as the engineering. The hope is that it will be possible to design drives that are *a priori* prone to escape many forms of resistance evolution. That is our thesis here. We expand on earlier work showing it was possible to develop moderate-effect suppression drives that would ‘out-run’ unlinked mechanism-blocking resistance (type-M) already existing in the population (Gomulkiewicz et al. 2021). But there were limits on how much population suppression could be achieved while evading resistance. We find here that ways exist to allow even greater magnitudes of population suppression while evading resistance evolution. Critically, this resistance must be at least partly unlinked to the drive, because the drive allele cannot otherwise fix (deterministically). However, we also show that this principle for evading type-M resistance does not translate equally to other resistance types. We also explore resistance evolution in the context of ecology.

With type-M resistance against a homing drive, there is the possibility of non-functional resistance – target-site mutations that block the drive and destroy the target-site gene function. Non-functional resistance can affect the spread and long-term impact (Beaghton et al. 2019). Non-functional resistance will be ignored here.

Our study assumes that resistance is unlinked to the drive. This assumption is not typical of other studies – most analyses of resistance to a gene drive have assumed that resistance is allelic to the drive (e.g., Unckless et al. 2015, 2017; Beaghton et al. 2017; Noble et al. 2017; Prowse et al. 2017; KaramiNejadRanjbar et al. 2018), although for an exception see (Charlesworth and Hartl 1978). Allelic resistance is highly relevant to a homing endonuclease for which resistance may take the form of a mutation in the target site (Champer et al. 2017; Noble et al. 2017; Bier 2021; Hammond et al. 2021; Terradas et al. 2021) as well as to some toxin-antidote systems, but there are designs that may avoid allelic resistance (see Discussion). Many other forms of resistance will not be allelic to the drive allele and not involve mutations at the cleavage site, such as blocks to gene drive expression, interference with endonuclease protein complexes, inbreeding, and others. The evolution of non-allelic resistance is fundamentally different from allelic resistance because the benefit of resistance no longer accrues directly to the resistance allele. The genetic separation of drive and resistance weakens selection of resistance and allows some forms of suppression drives to escape resistance evolution (Gomulkiewicz et al. 2021).

We address avoidance of resistance evolution in several diverse contexts. The cost of such diversity is that each case can be explored in only some of the vast diversity of possible variations that are of interest: drive imperfection, fitness effects of different genotypes, dominance/recessivity within loci, deviations from random mating, and ecology. To keep the study manageable, most of our models assume perfect drive with fitness effects in only the drive homozygote and no fitness cost to resistance: these are extremes that represent best cases for the evolution of resistance (no fitness cost) with a drive that has the best chance of outrunning resistance. The present study builds on our previous study that explored some of these variations (Gomulkiewicz et al. 2021). The application of principles found here to any implementation will require models specific to that case.

## Models and Results

Our methods use mostly numerical analyses of deterministic, discrete-time gene frequency models; the ecology section extends the models to population dynamics (code access is addressed in the Data availability statement). Mating is random unless specified otherwise. Two or three unlinked loci are assumed, each with 2 alleles. Some models assume that the sexes are haploids, other models assume diploid sexes. Computation employed C programs with graphics in R (R Core Team 2019).

### Two homing drives can escape resistance while achieving greater total suppression than does a single drive: 1-sex drives

It was previously shown that moderate-effect suppression drives could evolve to fixation even when resistance was initially present. The drive’s fitness effect was fully recessive, manifest only in drive homozygotes, and segregation distortion was 100%. Furthermore, and unexpected, suppression drives could evade resistance even if there was no fitness cost to the resistance allele (Gomulkiewicz et al. 2021); those models assumed type-M resistance.

In that prior study, there were limits on the suppressive effect of the drive that could evade resistance. The maximum possible suppression decreased with linkage between the resistance and drive loci, decreased with initial frequency of the resistance allele, but the possible suppression was considerably increased if the drive’s fitness effect was manifest in only one sex (Gomulkiewicz et al. 2021). Two-sex drives were only slightly better at evading cost-free resistance than one-sex drives when drive fitness effects were experienced in both sexes. (To clarify, ‘one-sex’ and ‘two-sex’ drives refer to the drive’s selfish advantage operating in one sex, such as only males, or both sexes.) Here we report that the fitness suppression attainable from ‘resistance-proof’ drives can be substantially improved when introducing 2 drives whose fitness effects combine multiplicatively. The introduction of multiple, simultaneous suppression drives was in fact proposed by Burt (2003) as a means to force a population to low numbers before resistance could ascend; our emphasis is somewhat different in that we consider drives that can fix in the presence of resistance. We confine this part of our analysis to type-M resistance and drives that operate in males only (one-sex drives), as that is a slightly more difficult case for evading resistance. Male drives have the advantage of avoiding Cas9 carryover to and unwanted nuclease activity in the embryo (Champer et al. 2017).

The model has 3 unlinked loci, each with 2 alleles: *A/a* (homing drive 1), *B/b* (homing drive 2), and *R/r* (resistance), lowercase alleles representing wild-type; resistance acts equally against both drives. Other assumptions about segregation distortion and fitness effects are provided in Tables 1 and 2. Complete segregation distortion operates independently at the *A/a* and *B/b* loci; dominant, cost-free resistance to segregation distortion operates at the *R/r* locus (Table 1).

**Table 1.**
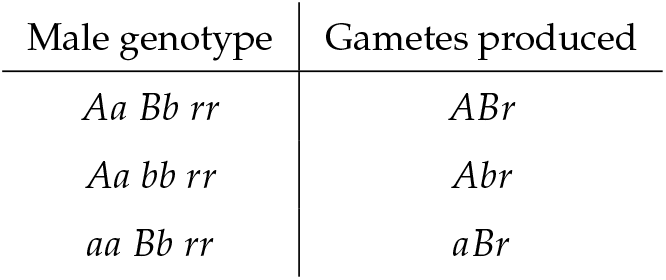
Gamete production for the three genotypes that are heterozygous at drive loci and wild-type homozygous at the resitance locus. Drive operates in males only via homing. Segregation is Mendelian for the 24 other possible male diploid genotypes not shown here and for all 27 possible female diploid genotypes. Resistance to segregation distortion is complete and dominant, and the *R* resistance allele fully blocks drive by *A* and *B*, separately and together.

| Male genotype | Gametes produced |
| --- | --- |
| *Aa Bb rr* | *ABr* |
| *Aa bb rr* | *Abr* |
| *aa Bb rr* | *aBr* |

**Table 2.** Diploid genotype fitnesses for the 3-locus model with two homing drives and a resistance locus. A dash (–) indicates either allele at the locus. Drive alleles are *A* and *B*, resistance is *R*. The parameter *s* determines the fitness reduction in drive homozygotes; fitnesses are multiplicative over loci. All genotypes not shown have fitness 1. Fitness effects operate on adults during gamete production. In some models, the fitness effects apply to both sexes, in others just to females, as given in Fig. 1. Cost-free resistance is assumed, as it is the most favorable case for resistance evolution. If a drive can evade cost-free resistance, then it should also evolve in all cases of costly resistance. Note that fitness of a genotype translates into population fitness to the extent that the genoytpe comprises the population.

| Genotype | Fitness |
| --- | --- |
| *aa bb rr* | 1 |
| *AA b- --* | $\sqrt{1-s}$ |
| *a- BB --* | $\sqrt{1-s}$ |
| *AA BB --* | $1-s$ |

Figure 1 compares the maximum *s* attainable by one drive versus 2 drives while evading cost-free resistance. These curves apply to drive alleles that evolve to fixation, after which type-M resistance can no longer influence evolution or fitness. The extreme case of resistance-free suppression (70%) is attained using 2 drives with equal fitness effects confined to just females. These figures assume that the two drives are introduced sequentially (second drive introduced when first drive has attained a frequency of 0.9995, dashed curves), but essentially the same curves are obtained for simultaneous introduction, suggesting that the major benefit of two drives is not from some interaction between the combination of two drive alleles and the resistance allele.

**Figure 1.**
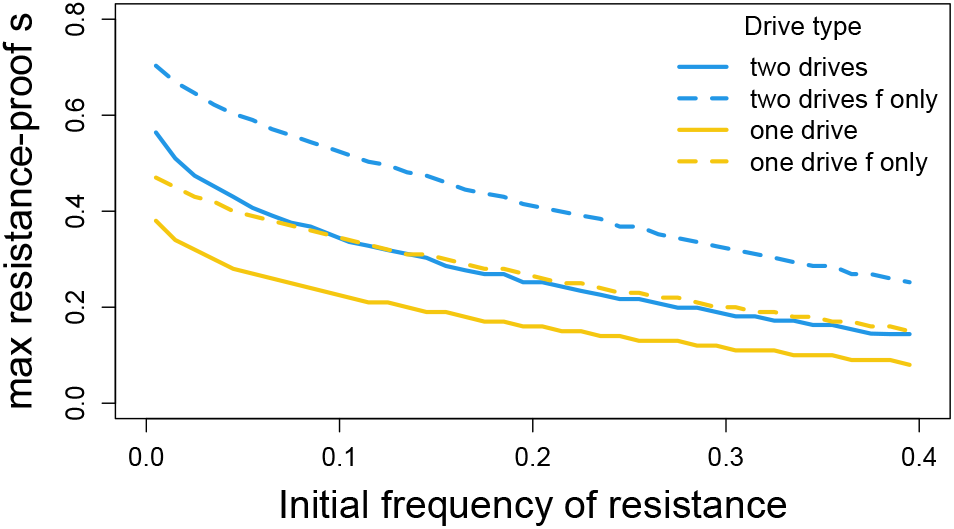
Maximum fitness suppression attainable with 1 versus 2 homing drives evolving in the presence of strong, cost-free resistance alleles; drive operates in males only. Curve height is the maximum fitness suppression attainable (*s*). When two drives are introduced, *s* is the combined effect of both drives, from Table 2. If only a single drive is introduced, *s* is the effect of the one drive (its homozygote fitness is 1 − *s*). For each initial frequency of resistance, the maximum resistance-free *s* was obtained over 100 trials spanning different levels of gene drive fitness effects (incremented every 0.01). The main point of the graph is that two suppression drives (blue) can escape resistance evolution and attain greater combined population suppression than can a single drive (yellow) of the same total effect; the advantage of 2 drives applies whether the fitness effects are experienced by both sexes (solid blue versus solid yellow) or just females (dashed blue versus dashed yellow), but a larger *s* can evolve resistance-free if the fitness effect is confined to females (dashed versus solid of same color). There is essentially no difference whether the two drives are introduced concurrently or sequentially, so sequential drive outcomes are shown. These suppression maxima were enduring, the drive allele(s) having fixed despite resistance being present in the population. The resistance allele persists but has no effect once the drive is fixed. The horizontal axis is the initial frequency of the (dominant) resistance allele, and it is easily seen that greater resistance-free suppression depends on a lower starting frequency of the resistance allele. Initial frequencies of the drive alleles were 0.005 when two drives were present, 0.01 for a single drive; some trials used initial frequencies of 0.0005, with no appreciable change from results with initial frequencies of 0.005. The leftmost initial frequency of the resistance allele was 0.005, incremented by 0.01.

The greater impact of resistance-proof suppression by using 2 drives instead of a single drive begs some understanding. We addressed this question with numerical results using our observation that 2 sequential drives can attain approximately the same total suppression as 2 concurrent drives; properties of gene frequency evolution were analyzed during single drive fixation (confining the analyses to parameter values for which the drives evolved to fixation). We found that two outcomes work in favor of greater resistance-free suppression by 2 weak drives than by 1 stronger drive. (1) For the same initial frequency of the resistance allele, larger values of *s* disproportionately lead to higher final frequencies of the resistance allele. (2) For the same value of *s*, higher initial frequencies of the resistance allele yield higher gains of the resistance allele (only noticeable at larger values of *s*). A third outcome works against resistance-free suppression by 2 drives: to attain a combined suppression of 1-s, the selective coefficient per drive needed when using 2 drives is slightly more than s/2, the excess increasing with s. The relative gain from using 2 drives is thus due to the net balance of these three components. A theoretical possibility is that arbitrary resistance-free population suppression could be obtained by using many drives of very small effect, but the method is not obviously practical beyond two or three drives.

#### Robustness

The model assumed a few extremes: perfect homing drives and no fitness costs to drive heterozygotes. To address the importance of the first extreme, trials were analyzed with drive distortion of 0.9 (drive heterozygotes produce 90% drive and 10% wild type gametes); this analysis was conducted for two male drives introduced simultaneously with equal fitness effects in both sexes and compared to that of a single male drive (corresponding to the solid lines in Fig. 1). There was little effect of this level of drive imperfection: the difference in maximum resistance-free *s* between a single drive and two drives changed less than 0.01 when comparing perfect drives to imperfect drives. A second check on robustness was to introduce a fitness decrement in drive heterozygotes. Surprisingly, a reduction in heterozygote fitness of 0.1 (from 1.0 to 0.9) increased the maximum resistance-free *s*. For perfect drive, the gain in resistance-free *s* was substantial even for a single drive (e.g., 0.06) and often somewhat larger for two drives (up to 0.09). For imperfect drives (90%, 10%, as above), the gain in resistance-free *s* by a 0.1 reduction in heterozygote fitness could be somewhat larger or smaller than the gain for perfect drives. The main observation remains that the use of 2 drives can attain a larger net suppression than the use of a single drive despite moderate variations in drive perfection and heterozygote fitness. It is of course understood that drive properties differing between the 2-drive and 1-drive cases could affect the benefit of using two drives.

### Inbreeding can evolve to block homing drives of even modest effect: 2-sex drives

The models both above and in previous work (Gomulkiewicz et al. 2021) assumed type-M resistance, thus blocking segregation distortion in the individual heterozygous for the drive(s). Resistance evolves because of a statistical association between the resistance allele and the drive-sensitive allele(s). Type-P resistance (population structuring) offers a different way to develop a statistical association between drive-sensitive alleles and resistance. We thus consider whether type-P resistance behaves similarly as type-M resistance in allowing modest-effect drives to fix without selecting resistance.

To represent type-P resistance, we invoke sib mating (Bull et al. 2019). Sib mating is not resistance in the sense of blocking segregation distortion, indeed it has no effect on distortion. Rather it provides a population structure in which drives are disproportionately partitioned across groups of individuals, thereby increasing the variance in fitness across families. In addition, the inbreeding itself reduces the frequency of drive heterozygotes, which are the only genotypes that permit the drive to collect its ‘unfair’ transmission advantage. In contrast to previous sib-mating models (Bull et al. 2019), we here allow drive homozygotes to be partly viable – a necessary condition for type-M drives to evade low levels of pre-existing resistance and thus an appropriate parallel for comparison. To keep the math simple, we assume sexual haploids: mated females produce a brief diploid stage in which segregation distortion occurs if and only if the diploid is *Aa*, and fitness effects of drive occur if and only if the diploid is *AA* (Table 3). Since the diploid phase is neither male nor female, this model is effectively a 2-sex homing drive. There is no genetic resistance to the drive action as such, as all drive heterozygotes produce only *A* gametes (Table 3).

**Table 3.** Gamete production and fitnesses for the three drive genotypes in the sib-mating model. Drive operates in *Aa* diploids only and does not affect other loci. Drive is not suppressed or affected by other loci. These properties apply to all diploids. Diploids are not distinguished as male or female, as sexes apply to haploids. *s* is manifested as viability of *AA* progeny.

| Diploid genotype | Diploid fitness | Progeny produced |
| --- | --- | --- |
| $aa$ | 1 | $a$ |
| $Aa$ | 1 | $A$ (after drive) |
| $AA$ | $1 - s$ | $A$ |

Sib mating is enforced by the mother on her progeny (Table 4). Only mothers of genotype *Q* have sib-mated progeny, and then only a fraction *m* of those sib mate. All offspring of *q* mothers and 1 − *m* offspring of *Q* mothers join the random mating pool. We assume that males who mate with their sisters do not also join the random pool, which would give sib mating an advantage independent of gene drives (Lande and Schemske 1985). The fitness effect 1 − *s* is manifested as viability of *AA* progeny.

**Table 4.** Relationship between maternal genotype at the *Q/q* locus and the fraction of her progeny that engage in sib mating.

| Maternal genotype | sib mating fraction |
| --- | --- |
| $Q$ | $m$ |
| $q$ | 0 |

It is further assumed that sib mating does not affect offspring fitness (no inbreeding depression). This assumption is equivalent to cost-free type-M resistance and enables isolating the drive-blocking effect of sib mating. It is obviously a best-case scenario for the evolution of type-P resistance, and although it may be unrealistic, it should also define the conditions under which a suppression drive can be assured of evading sib mating.

Figure 2 shows three scenarios differing in the degree of sib mating imposed by mothers with allele *Q*, from low levels of sib mating (*m* = 0.2), medium (*m* = 0.5) to high levels (*m* = 0.95). Each graph displays equilibrium mean fitness (average female survival), equilibrium drive frequency, and equilibrium frequency of *Q* for different values of drive fitnesses (1 − *s*); (note that sib mating has no fitness consequences if and after the drive fixes, so polymorphism of *Q* after drive fixation merely reflects its spread prior to drive fixation). There are three important outcomes.

1. Sib mating does not evolve (and the drive evolves to fixation) unless the sib-mating fraction exceeds the fitness of the drive homozygote (*m* > 1 − *s*). Thus, mild-effect drives can avoid type-P resistance evolution if there are only low levels of sib mating types in the population.
2. When it evolves enough to at least partially suppress the drive, the sib-mating allele typically fixes. For some combinations, the drive allele can be maintained polymorphic despite fixation of allele *Q*. We have not observed joint polymorphism of *Q/q* and the drive allele, but we cannot rule it out.
3. If the sib-mating genotype enforces a high degree of sib mating (*m* near 1), sib mating evolves to extinguish all but the most benign gene drives.

**Figure 2.**
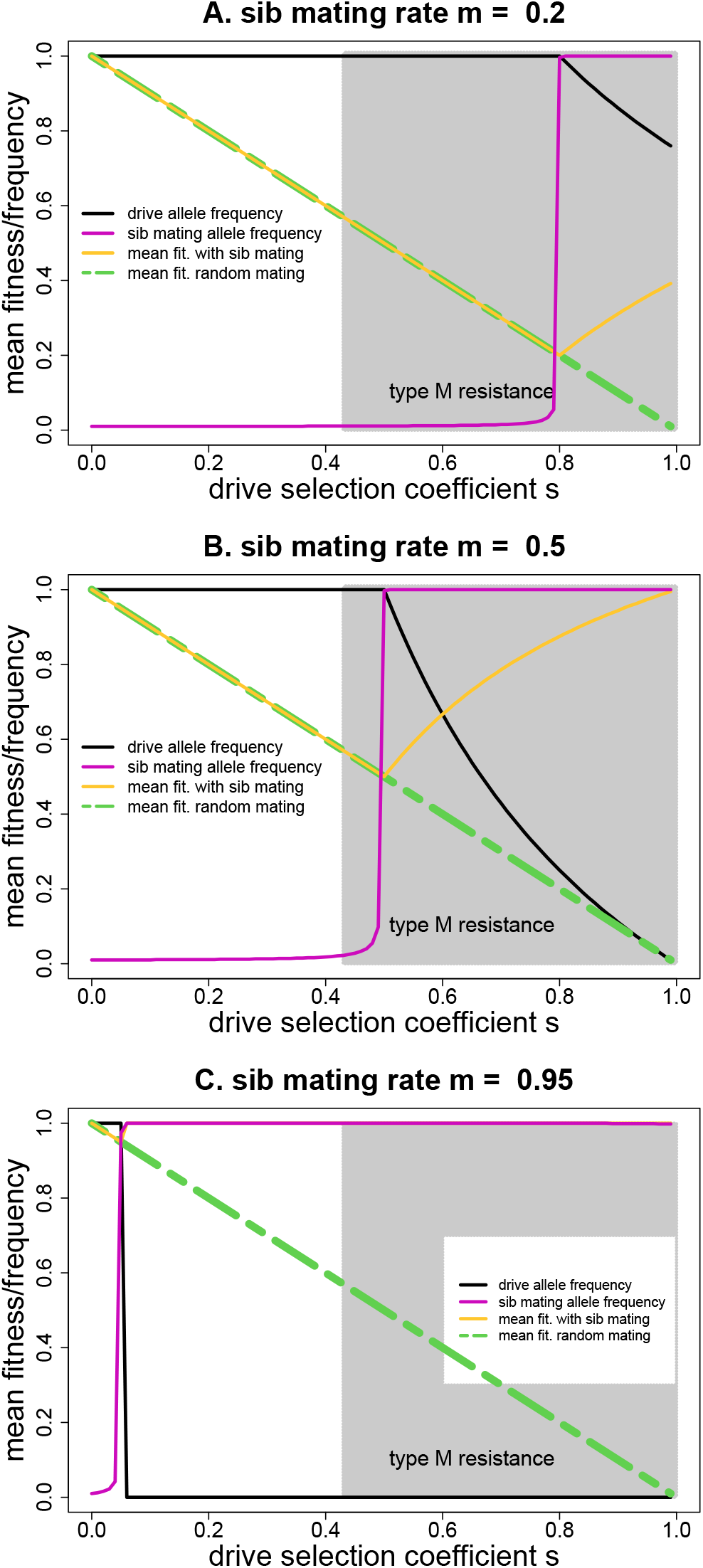
Equilibrium consequences of homing drive evolution when confronted with evolution of population structure resistance (type-P) in the form of sib mating. The black curve provides the equilibrium frequency of the drive allele. The orange curve gives evolved mean fitness (average female survival, since male frequencies were normalized to 1); the green curve gives equilibrium fitness in the absence of sib mating; the pink curve gives the frequency of the sib-mating allele *Q*. Panels differ in the level of sib mating, *m*, imposed by the sib-mating allele *Q*: 0.2 (panel A), 0.5 (panel B), 0.95 (panel C). Analytical results (supplemental file S1) show that the gene drive fixes if selection against it is below the level of outcrossing enforced by the inbreeding allele (i.e., if *s* < 1 − *m*) and is lost if *s* > 1/*m* − 1; at extremes of *m*, the drive allele is vanquished. The numerical results agree with that behavior. For 1 − *m* < *s* < min(1/*m* − 1, 1), polymorphic drive equilibria exist with allele frequency equal to 1/*s* − *m*/(1 − *m*). Overall, *Q* alleles with high values of *m* evolve to fixation at all but low values of *s*, but whatever the value of *m*(> 0), *Q* will evolve to fixation as *s* gets high enough. But once a value of *s* is reached such that *Q* fixes, if the drive allele remains polymorphic, increased values of *s* lead to lower drive frequency and higher mean fitness. For comparison, the shaded area shows the values of *s* for which an unlinked, type-M resistance allele would evolve to block the drive. The curves in each panel were generated from trials incremented every 0.01 on the X-axis; initial frequencies were 0.01 for both the sib-mating allele and the drive allele (or the resistance allele for type-M resistance). Outcomes were assessed at 10^5^ generations. Note that in (C), the orange curve follows the green only at low values of *s* and then rises to join the pink throughout most of the range of *s*.

In comparison to homing drive evasion of type-M resistance, sib mating provides a mixed message. Modest effect drives can fix if the population lacks alleles that enforce high levels of sib mating, even if the population has alleles that enforce low levels of sib mating (Fig. 2A, B). This offers a parallel to modest-effect drives evading type-M resistance whereas lethal drives cannot evade resistance. In contrast, a recessive lethal drive (*s* = 1) will select any level of sib mating (all *m* >0), although the equilibrium frequency of the drive remains high if the sib mating allele enforces only low levels of sib mating (Fig. 2A).

Seemingly without parallel in type-M resistance, homing drives of moderate or even small effect cannot evolve to fixation if the population has genotypes that enforce high levels of sib mating (Fig. 2C). In this sense, sib mating provides a graded resistance response – if the population has even a low frequency of genotypes that impose high levels of sib mating, they will evolve to stop even weak-effect drives. But if the population has only genotypes that impose low levels of sib mating, even moderate-effect drives can evolve to fixation. Unfortunately, it may be difficult to anticipate in a wild population what levels of genetically controlled sib mating exist at low frequency.

There is an interesting difference in drive evolution between the *m* = 0.95 case and the other two cases considered in Fig. 2. When the *m* = 0.95 sib mating allele evolves to fixation, the drive allele is purged. In contrast, for *m* = 0.2 and *m* = 0.5, the drive remains polymorphic or even abundant despite fixation of the sib-mating allele. Persistent drive polymorphism could select additional types of resistance.

#### Robustness

These results were based on perfect drive and no heterozygote fitness effect of the drive. Revisiting the three cases in Fig. 2, those assumptions were relaxed by allowing a moderately imperfect drive (drive heterozygotes produce 90% drive and 10% wild type gametes) and/or a heterozygote fitness cost of 0.1. The effects were quantitative and did not alter the overall outcomes. Drive imperfection reduced the range of *s* values in which drive fixed by 0.01 for *m* = 0.95 to 0.05 for *m* = 0.2. Mean fitness in the zone of drive polymorphism increased substantially with drive imperfection alone for *m* = 0.5 and *m* = 0.2 (by as much as 0.18 and 0.26, respectively), with the heterozygote fitness cost also increasing mean fitness but less so. The basic result that high magnitudes of sib mating purge drives of even small *s* effect remains. The fact that drive imperfection reduces population suppression in the zones of drive polymorphism is not surprising, and the magnitude of effect may have little permanance given that the polymorphic drive would be prone to select other forms of resistance.

### Sib mating can evolve in response to toxin-antidote drives, but patterns differ from homing drives

Inbreeding has a moderately straightforward interaction with homing drives, which are confined to one locus. If the principle behind the evolution of sib mating in response to a suppression drive is that it increases the variance in fitness across families, one would expect sib mating to be favored with other forms of suppression drives. Here we consider evolution of sib mating in response to toxin-antidote systems used for suppression.

There is a large variety of possible toxin-antidote designs (Champer et al. 2020a). Our example of a toxin-antidote drive is ClvR (Oberhofer et al. 2019) or equivalently distant-site TARE (Champer et al. 2020b). Two loci are involved. At the ClvR locus, with alleles *c/C*, the *C* allele encodes both a rescue gene and a nuclease. The nuclease annihilates wild-type alleles at a second, unlinked, essential-gene locus, *g/G* (Table 5). ClvR (allele *C*) is engineered as a one-locus construct with two functions, the *g/G* locus merely being the passive ‘victim’ of the nuclease of the first locus. The *C* allele spreads because it creates non-functional *G* alleles, with *GG* genotypes dying if not rescued by *C* (*ccGG* is the only genotype that dies because it lacks the rescuing *C* allele). ClvR does not intrinsically suppress populations. To be used for population suppression, the *C* can be inserted into a gene important to survival/fecundity so that the gene is disrupted and homozygotes impaired (Table 5).

**Table 5.** Genotypes, fitnesses and functions of the 2 loci in the ClvR toxin-antidote system; a dash (-) represents either allele. Allele *C* performs two functions: it converts all wild-type *g* alleles in the individual to non-functional *G*, and it rescues *GG* from death. The fitness parameter here is *σ*, to distinguish it from *s* in homing drive models. These properties apply to all diploids. Diploids are not distinguished as male or female, as sexes apply to haploids. Assigning a fitness decrement to *Cc* heterozygotes imposes a threshold frequency requirement for spread of *C* when rare (Oberhofer et al. 2019; Champer et al. 2020b).

| Genotype | fitness | behavior |
| --- | --- | --- |
| $cc\ g-$ | 1 | |
| $cc\ GG$ | 0 | |
| $Cc\ g-$ | 1 | $g \rightarrow G$ |
| $Cc\ GG$ | 1 | |
| $CC\ g-$ | $1-\sigma$ | $g \rightarrow G$ |
| $CC\ GG$ | $1-\sigma$ | |

When *CC* fitness is impaired, ClvR faces the same threat of allelic resistance as do homing drives. A target-site allele *g* that is resistant to cleavage but retains function will eventually ascend and prevent evolution of *C*. The same designs used for avoidance of functional allelic resistance in homing drives will thus need to be used for ClvR (see Discussion). Our interest here is in the evolution of sib mating as a form of type-P resistance. Sib mating operates as in the previous model (Table 4); the model again assumes haploid sexes, but fitness effects and allelic conversion occurs in diploids (Table 5).

In contemplating the evolution of inbreeding, use of ClvR for population suppression exhibits some notable differences from homing drives. Note that we are confining our analysis to viability effects of *C* on both sexes:

- In the absence of resistance, evolution of ClvR leads to a population that is fixed for null homozygotes *GG* but polymorphic for *C/c*: *CC* has low fitness, *Cc* is normal, and *cc* dies because all wild-type *g* are gone. This fitness overdominance holds for any *σ* > 0 and is a basis for inbreeding depression. The equilibrium frequency of C with a fixed inbreeding coefficient *f* is 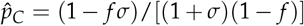. At least when rare, inbreeding would likely be selected against as the population neared this state since less inbred families would produce more descendants. This behavior contrasts with that of homing drives considered above, which have no effect on sib mating once they are fixed.
- Inbreeding affects and is affected by two loci with ClvR: the *c/C* locus (as noted above) and the *g/G* locus, for which GG is lethal in the absence of *C*. With homing drives, only one locus is affected by inbreeding.
- As elaborated in the next section, ClvR cannot suppress mean fitness below 0.5 (when fitness effects are confined to the homozygote as viability effects). The effect of fitness parameter *σ* on mean fitness is not the same as that of the homing drive parameter *s*.

These differences raise the possibility that sib mating will evolve differently under ClvR than under homing drives.

Plots of sib mating evolution in response to ClvR evolution show similar patterns as for homing drives in that strong allelic sib mating can evolve to block the suppressing ClvR (compare Fig. 3 to 2). There are two differences worth noting, however:

1. Alleles enforcing weak sib mating evolve only at higher values of *σ* than for *s* with homing drives. For example, with *m* = 0.5, ClvR does not select sib mating until *σ* is almost 0.7, whereas homing drives selected sib mating when *s* reached 0.5.
2. At equilibrium, joint polymorphism of *C* and sib mating was never observed, although we cannot rule out the possibility. If sib mating evolved, ClvR was lost, at which point the *Q* allele was neutral. Yet with homing drives, the drive allele could be maintained after *Q* evolved to fixation (for some conditions).

**Figure 3.**
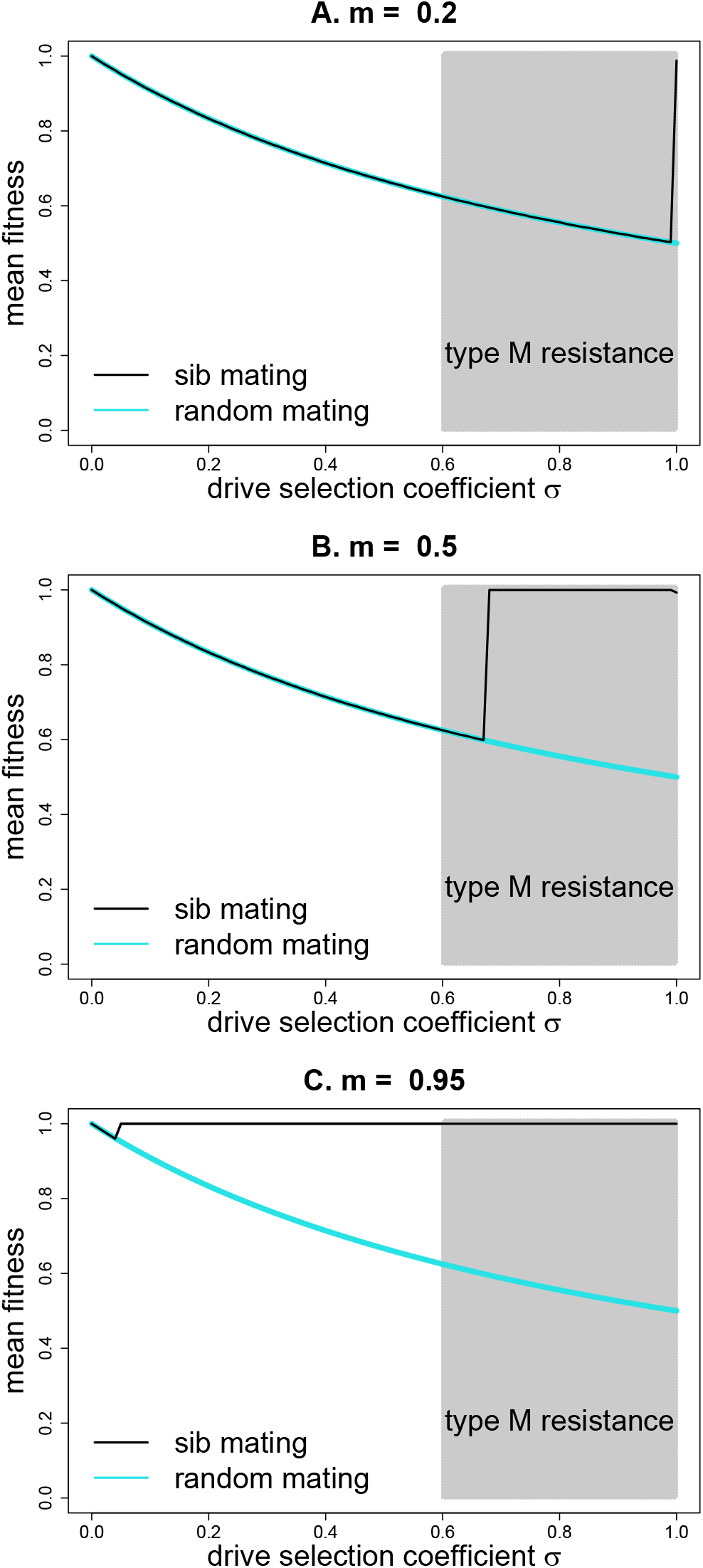
Equilibrium mean fitness (average survival) for toxin-antidote (ClvR) evolution when confronted with evolution of sib mating. Panels differ in the level *m* of sib mating imposed by the inbreeding allele: 0.2 (top panel), 0.5 (middle panel), 0.95 (bottom panel). The blue curve represents equilibrium mean fitness when ClvR evolves without interference from sib mating, 1/(1 + *σ*). Sib mating evolves to suppress the drive where the black curve rises above the blue; the black curve represents evolved mean fitness. In contrast to sib mating suppression of homing drives, the effect on ClvR is close to all or none – mean fitness is virtually always 1.0 or that expected if ClvR is unaffected. The shaded area spans the values of *s* for which type-M resistance evolves in the same model (even though *s* and *σ* have somewhat different effects). As with homing drives, sib mating can evolve at much smaller values of *σ* if the sib mating allele enforces a high level of sib mating. Initial frequencies of the ClvR and sib-mating alleles were 0.07; initial frequencies in the type-M trials were 0.01. Trials were evaluated at 20,000 generations.

These results may not be representative of all outcomes at least because our explorations are limited to one set of initial frequencies. We expect that the interior dynamics of the joint evolution of ClvR and sib mating to be complicated because of the interaction among the 3 loci. Indeed, cursory numerical explorations have observed internal equilibria. The main point is that, as with suppression homing drives, a suppression toxin-antidote system can select inbreeding under a wider range of *s* than for which type-M resistance evolves.

### Fitness suppression potential across different drive systems

The goal of a suppression drive application is to reduce actual population sizes of a species. Analyses above were based on average fitness effects, a genetic measure, and the translation between a reduction in fitness and the demographic effect on population size can be complicated. This section offers a prelude to the study of ecological effects by revisiting how mean fitness reduction depends on the type of drive (Fig 4).

**Figure 4.**
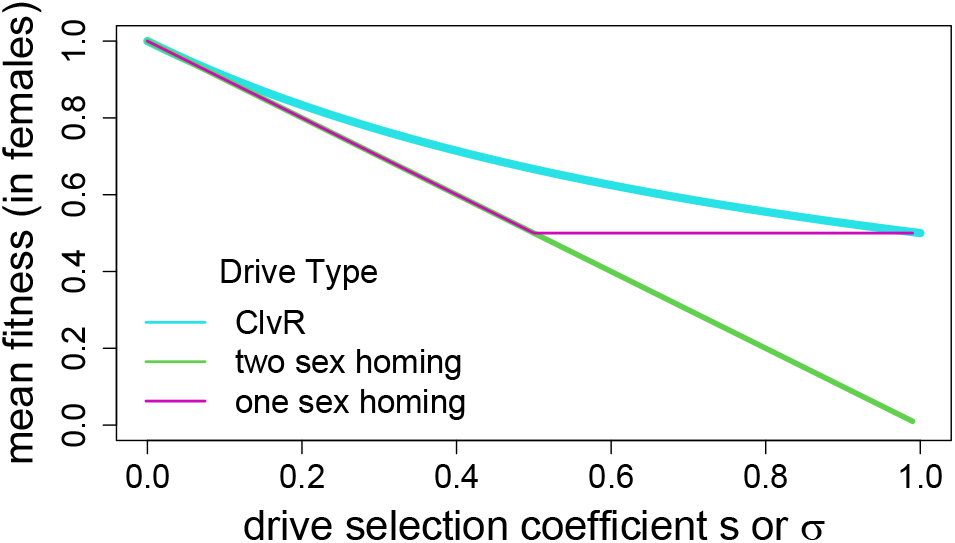
Equilibrium mean fitness (average survival) as a function of the drive homozygote fitness effect for single 1-sex homing drives (purple), 2-sex homing drives (green), and the ClvR toxin-antidote drive (blue); *s* applies to homing drives, *σ* to ClvR. These outcomes assume that the fitness effect operates in both sexes, that drive has evolved to its maximum level, and that resistance is absent; similar curves apply if the fitness effect is confined to one sex and mean fitness is measured in that sex. Only 2-sex homing drives can suppress mean fitness below 0.5 for these systems, although some toxin-antidote systems can suppress mean fitness even lower (Champer et al. 2020a). A comparison of fitness effects across different drive systems is also provided by Beaghton et al. (2019).

In the absence of resistance, a 2-sex homing drive can evolve to suppress mean fitness anywhere from 1.0 to 0 depending on the fitness 1 − *s* of the drive homozygote (see Prout 1953, for the case of *s* = 1). In contrast, a 1-sex homing drive can suppress mean fitness no lower than 0.5 (Bruck 1957). When *s* exceeds 0.5 in a 1-sex homing drive, the equilibrium population is polymorphic for the drive because the non-drive sex produces 50% wild-type gametes; mean fitness remains at 0.5 for the entire range of 0.5 < *s* < 1.

ClvR, our representative toxin-antidote system, has elements of 2-sex and 1-sex homing drives. It operates in both sexes, but it’s equilibrium fitness mirrors that of 1-sex homing drives. Thus, as a consequence of *c/C* polymorphism at equilibrium (noted above), the most extreme fitness suppression occurs when both homozygotes die, leaving *Cc* heterozygotes as the only viable genotype and a mean fitness of 0.5 (Fig. 4).

The figure shows that 2-sex homing drives can achieve far greater fitness suppression than either 1-sex homing drives or ClvR. Perhaps surprisingly, there is little difference between 1-sex and 2-sex drives much less difference in the resistance-free zone of fitness suppression (Gomulkiewicz et al. 2021). In the preceding sections of this paper, our sib-mating models were effectively 2-sex drives, but the section comparing two drives with one drive assumed a 1-sex drive. The conclusions of those sections should not be qualitatively sensitive to the assumptions of 1-sex versus 2-sex drives.

The next section delves further into ecology but continues with the theme of resistance evolution. It is limited to homing drives, but the principles should apply to toxin-antidote drives as well.

### Avoiding resistance using ecology: homing drives

Many applications of gene drives will be designed for ecological effects – population suppression or eradication. It is not straight-forward to translate the relative fitness effect of a gene drive into ecology: how does a 50% reduction in birth rate translate into a change in adult population size, for example? Furthermore, it may be possible to exploit desired ecological effects while evading evolution of resistance by choosing ecological traits that amplify effects on population size while reducing relative fitness effects. One such example is to limit drive fitness consequences to females, thereby essentially halving the relative fitness effect but yielding the same impact on population growth as if males were also affected.

Although ecological processes collectively offer a one-to-many translation of relative fitness into adult population size, depending on assumptions about ecology (e.g., Nagylaki 1992), we expect that the type of drive will not strongly affect this translation. Thus, we expect that a drive reducing mean fitness to 0.5 will have largely the same population impact regardless of whether the drive is a 1- or 2-sex homing drive or a toxin-antidote drive (assuming no resistance). There will be differences in the speed of evolution, and speed could affect the evolutionary response and/or the ecology.

There are countless possibilities to consider at the interface of gene drives and ecology. Our focus continues to be that of avoiding resistance evolution. We develop a model to address the relationship between population genetics of gene drives, resistance evolution, and ecological impact on population. The model is deterministic and assumes discrete generations, random mating of diploid males and females, with drive operating only in (heterozygous) males. To model population size, the number of progeny born is determined by the number of adult females; males are never limiting for female reproduction. Birth rates and viability effects on females therefore determine population size in each generation based on numbers of adult females in the previous generation.

Fitness effects of the drive affect progeny viability regardless of sex and may operate in a density-independent manner as well as a density-dependent manner (Table 6). The density-dependent effects are analogous to effects on ecological carrying capacity (though not strictly the same). Following previous sections of this paper, our basic question is how gene drive avoidance of resistance depends on an ecological context. Can the ecology be used to further evade resistance?

**Table 6.** Genotype-specific birth rate and progeny viability in the homing drive model of ecological effects. A mother’s number of surviving offspring is the product of her births (*b*, genotype-independent) and the survival of her offspring according to their genotypes. The net survival of each progeny is the product of a density-independent and density-dependent term (given in the third and fourth columns) and depends on their genotypes. The density-dependent model is a type known as ‘ceiling’ density dependence (Lande 1993). *N* is the total number of progeny born in the population that generation, prior to viability effects. *K* is the threshold density of *N* for the influence of density-dependent effects. A birth rate term is included here to emphasize that the birth rate affects the population dynamics and evolution even though there is no genotypic effect on birth rate. These rules used for enforcing density dependence are convenient for numerical analysis, and they have the advantage of preventing major fluctuations in adult population sizes that can occur with many other types of density dependence.

| drive<br>genotype | mother's<br>birth rate | density-ind.<br>viability | density-dep.<br>viability |
| --- | --- | --- | --- |
| $a-$ | $b$ | 1 | $\min(1, \frac{K}{N})$ |
| $AA$ | $b$ | $1 - s_b$ | $\min(1, \frac{K(1-s_K)}{N})$ |

Much of the evolution in this model can be qualitatively anticipated by translating the information of Table 6 into density-dependent effects on just the drive homozygote, *AA*. Its density-dependent viability descends from 1 to 1 − *s_K_* across three zones determined by the population size *N* relative to the thresholds *K* and *K*(1 − *s_K_*) (Table 7). Comparing density-dependent to density-independent viability (for *s_b_* = *s_K_*), density-dependent selection against *AA* is never more extreme than density-independent selection and may be considerably less. Whenever *N* is below the *AA* viability threshold of *K*(1 − *s_K_*), the drive has no relative fitness effect on *AA* – providing a ready means for ensuring resistance-free evolution by (for example) suppressing population size artificially.

**Table 7.** Relative fitnesses of *AA* genotypes as determined by offspring viability and according to the number of progeny born in the population in that generation (*N*); these relative viabilities follow directly from Table 6 but are presented here for clarity. There are 3 zones depending on the relative magnitudes of *N, K*, and *K*(1 − *s_K_*), and the zones apply to the density-dependent component. When the number of offspring born is below *K*(1 − *s_K_*), all genotypes (*aa, Aa*, and *AA*) have equal density-dependent fitness components; the relative fitness of *AA* declines as the number of offspring increases, but only down to 1 − *s_K_* – when the absolute viability of *AA* is *K*(1 − *s_K_*)/*N* but the absolute viabilities of *aa* and *Aa* are *K/N*. In contrast, the density-independent viability component is constant across all three zones.

| population size<br>(progeny) | relative<br>density-ind.<br>viability of $AA$ | relative<br>density-dep.<br>viability of $AA$ |
| --- | --- | --- |
| $N < K(1 - s_K)$ | $1 - s_b$ | 1 |
| $K(1 - s_K) \leq N \leq K$ | $1 - s_b$ | $\frac{K(1-s_K)}{N}$ |
| $K < N$ | $1 - s_b$ | $1 - s_K$ |

To study the impact of both types of viability effects on resistance-free evolution, numerical trials were conducted with different birth rates and with periodic depression of the population’s progeny; further model details are given in figure legends and tables. The clear result is that a homing drive with purely density-independent effects evades resistance according to the usual rules from population genetics models (Fig. 5, top). In contrast, a drive with purely density-dependent effects evolves differently; it evades resistance more easily when the population is frequently suppressed below its maximum. With the periodic population culling (knockdown) imposed by our model, resistance evasion is further enhanced by low birth rates, because low birth delays the population’s rebound to the viability threshold *K* (Fig. 5, bottom). As elaborated in the Discussion, the practical ramifications of this latter point are clear: evolution of resistance against a drive with a density-dependent fitness effect can be avoided by maintaining the population in a suppressed state that largely avoids the density-dependent effects.

**Figure 5.**
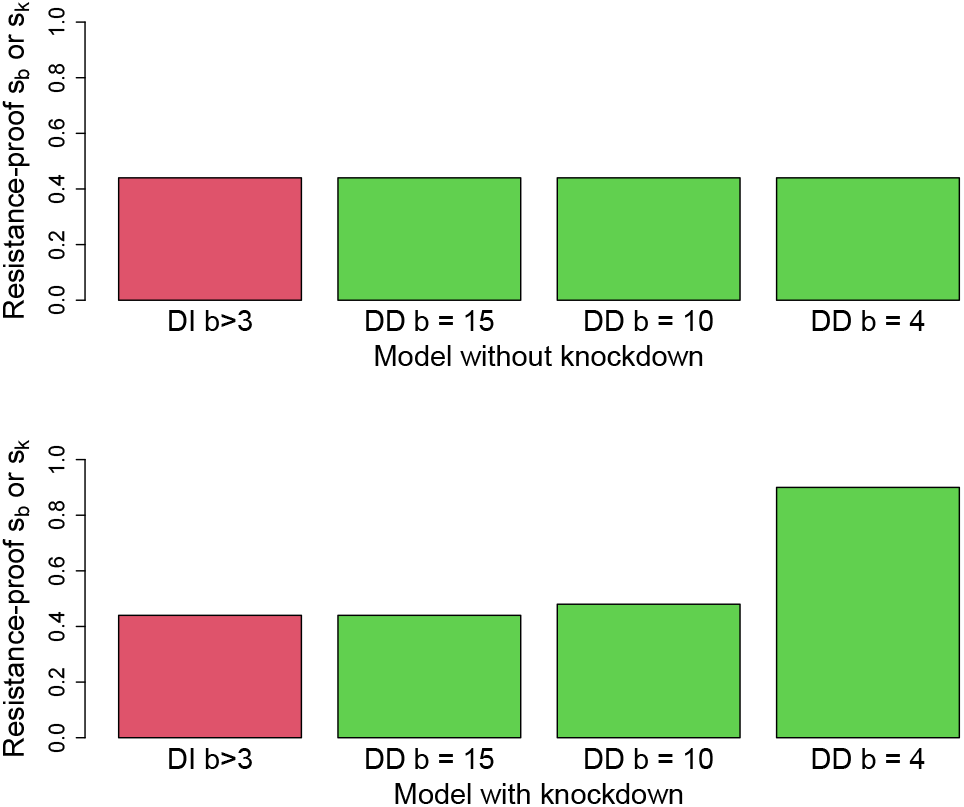
A drive that suppresses viability can evade resistance at higher fitness effects when its effect is density-dependent than when its effect is density-independent. Bars show the range of viability coefficients (*s_b_*, *s_K_*, Table 6) that allowed resistance-free evolution of a homing drive in the ecology model. The bars on the left of each panel (red, labeled with DI) apply to all trials in which the drive affected density-independent viability (*s_b_*); the three (green) bars on the right apply to trials in which the drive affected the density-dependent viability (*s_K_*), each specific to a different value of the intrinsic birth rate (*b*), as indicated. The DI bars combine many trials using different birth rates, as all resulted in the same resistance-free upper limit. (Top) Population numbers were determined solely by births and homing drive evolution, without any outside suppression or enhancement of numbers. All drives, whether affecting the density-independent or density-dependent viability parameter, showed the same resistance-free upper limit on evolution (approximately 0.43). (Bottom) Trials used the same conditions as in the top panel, except that total progeny numbers were artificially depressed every 5 generations to 1/4 of their initial values. Depending on the intrinsic birth rate, this depression could lower the total progeny (*N*) below the thresholds *K* and *K*(1 − *s_K_*) in Table 6. Furthermore, again depending on birth rates, the population could require several generations to recover before the next suppression. This suppression had no effect on the resistance-free upper limit when density-independent viability operated. However, with density-dependent viability effects, the range of resistance-free *s_K_* increased as *b* declined. This latter effect of *b* is expected because the temporary depression of population numbers is greater and the recovery slower with lower birth rates, hence the population experiences longer periods of reduced selection against the drive allele (as per Table 7) which in turn reduces selection for resistance (Gomulkiewicz et al. 2021). Parameter values and initial conditions were *K* = 1 × 10^9^, initial population size 1 × 10^9^, and initial frequencies of drive and resistance allele = 0.0005. The slight difference between the resistance-free zone of density-independent effects here and in Fig. 1 is due to the different initial frequencies used.

These results are so far limited to the evolution of resistance and then only showed final outcomes. The parameters were chosen so that none of the trials led to population extinction, but the evolution did sometimes result in suppressed densities. We now show how extinction can occur, finding that extinction can be achieved by a density-independent effect even when the density-dependent effect would allow persistence – all while avoiding resistance evolution.

From the above results, it is easy to appreciate that drive-caused extinction from a density-independent viability effect requires an intrinsically low birth rate. To avoid resistance, the drive cannot have too large of a viability reduction, *s_b_* (an approximate upper limit is 0.4 for these models, Fig. 5). Therefore, when the intrinsic birth rate *b* is near the minimum for population persistence, a density-independent *s_b_* = 0.4 can evolve resistance-free and suppress viability below that needed to maintain the population (Fig. 6); in contrast, if the intrinsic birth rate is high, this evolution will not lead to extinction. Perhaps surprisingly, under the very conditions that will cause extinction when the viability effect is density-independent, assigning the same viability effect to *s_K_* instead of *s_b_* will lead to lower adult population size, not extinction (Fig. 6). We suggest that this inability to cause extinction through density-dependent viability will be general in this (deterministic) model for any reasonable value of *K*. However, adding Allee effects or demographic stochasticity could lead to extinction with suppression of density-dependent viability (Lande et al. 2003).

**Figure 6.**
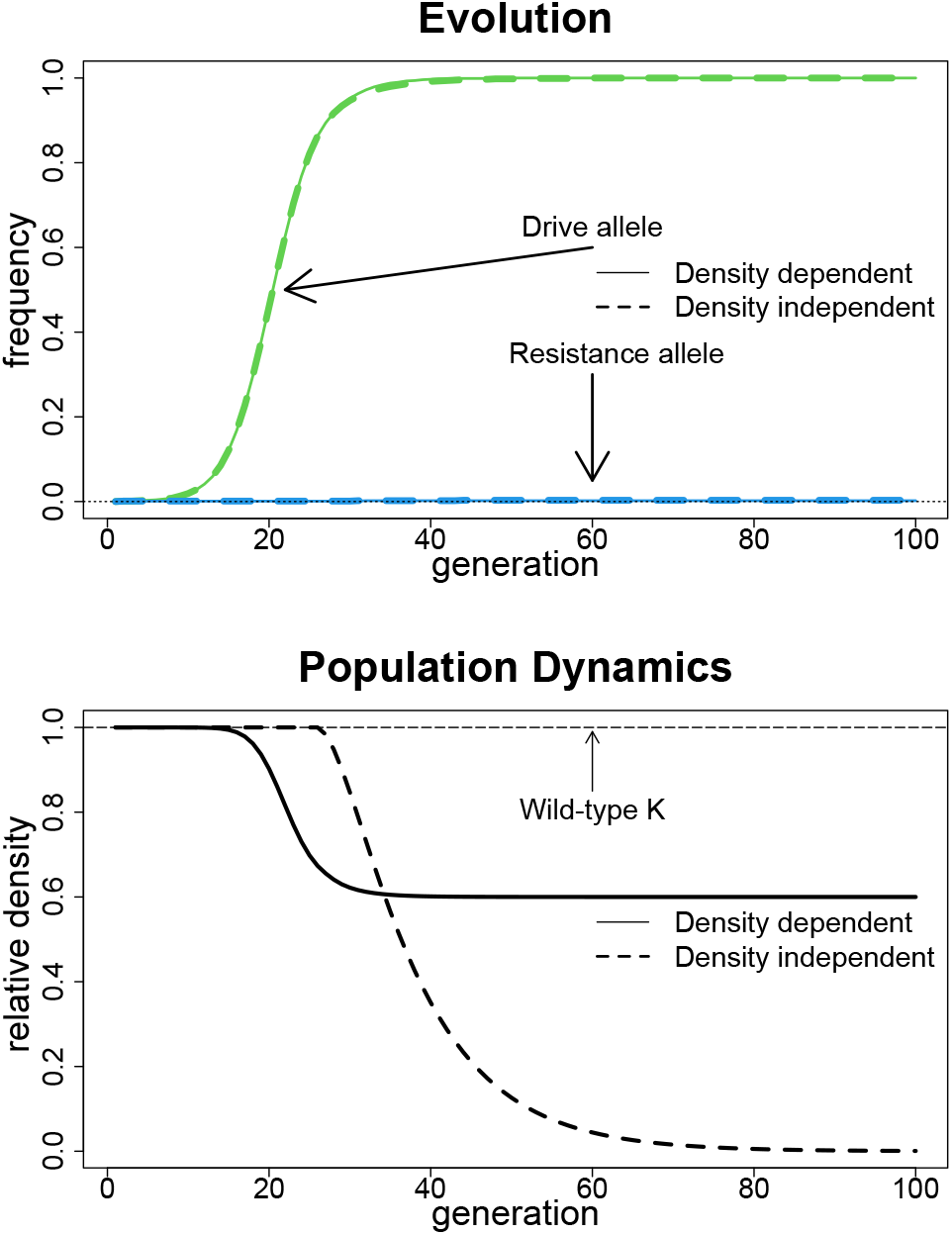
Drives with density-independent fitness effects can cause extinction when drives with the equivalent density-dependent fitness effects do not. Extinction requires a low intrinsic birth rate, so that the drive can evade resistance by having only a moderate fitness effect yet suppress births below that needed for population persistence. In contrast to other figures in this paper, the population dynamics over time are illustrated. The top panel shows the evolutionary dynamics of drive allele frequency (green) and of resistance allele frequency (blue) for the density-independent (dashed) and density-dependent (solid) cases. The evolution is approximately the same in both models. The bottom panel displays the dynamics of population densities (relative to *K*) for the two cases. Extinction occurs for the density-independent case as the density approaches zero, but not in the density-dependent case, for which the density declines by a factor 1 − *s_K_*. Parameter values are *b* = 3, *s_b_* = *s_K_* = 0.4, *K* = 1 × 10^9^. Initial allele frequencies were the same as in Fig. 5.

The ecology models developed here allow us to tentatively identify principles for resistance avoidance:

- Density-dependent fitness effects afford opportunities to relax resistance evolution. For example, drive-imposed lethality that operates only at high population density experiences reduced resistance evolution if the population is temporarily maintained at low density until the drive has swept. This population suppression can be manifest artificially or (in some cases) by the gene drive itself, the latter process being sensitive to intrinsic birth rates. Likewise, conditionally expressed fitness effects can avoid selection during drive spread and thus facilitate avoidance of resistance despite later manifestation of the fitness effects. For example, a drive that reverses pesticide resistance can evolve cost-free if pesticide use is averted until drive fixation.
- Density-independent fitness effects can achieve extinction while avoiding resistance evolution if intrinsic fecundity is low. There may be few pest species for which this is relevant though it may help with suppression of naturalized invasives.
- Fitness effects that operate in only one sex reduce resistance evolution while possibly enhancing or minimizing ecological effects. Thus, reducing only female fitness will more easily suppress populations than does reducing only male fitness, and fitness effects confined to one sex reduces selection for resistance (e.g., Fig. 1, and (Gomulkiewicz et al. 2021)).

## Discussion

At face value, the easy engineering of gene drives affords an ability to manipulate plant and animal pests that is unprecedented in the history of civilization. Merely releasing a handful of appropriately-engineered individuals of a species can potentially eradicate the entire species worldwide using suppression drives. This potential was known at least for the last half century (Hamilton 1967; Lyttle 1977), but engineering remained the hurdle. With engineering no longer a hurdle in gene drive construction, (e.g., Kyrou et al. 2018; Oberhofer et al. 2019; Champer et al. 2020a,c; Oberhofer et al. 2020; Simoni et al. 2020; Bier 2021), the unresolved question is whether resistance evolution will thwart the effort, as observed in some cage populations of the first experimental gene drive study (Lyttle 1979, 1981), but not invariably in more recent ones (Kyrou et al. 2018; Hammond et al. 2017). It may be impossible to accurately predict resistance evolution in any wild release (Burt 2003), since the outcome will depend on the nature of variation in the species, but modeling can at least help us to anticipate possibilities.

The conclusion here and in a previous study (Gomulkiewicz et al. 2021) is that engineering can substantially influence whether resistance evolves to block a suppression drive. Use of a highly suppressive drive will ensure evolution of resistance if resistance exists in the population or can arise quickly enough. Use of a moderately suppressive drive can escape resistance evolution. This paper elaborates on some of this understanding.

We distinguished three types of resistance evolution: resistance that blocks the drive mechanism (type M, Lyttle 1979; Champer et al. 2017), compensatory resistance that reverses the fitness effects of the drive without affecting drive (type C, as observed by (Lyttle 1981)), and resistance that affects population structure to prevent the drive from spreading (type P). We previously showed that suppression drives with moderate fitness effects could escape unlinked type-M resistance evolution even when the resistance alleles were present initially and imposed no cost on fitness (Gomulkiewicz et al. 2021). Resistance evolution depended on linkage disequilibrium between the resistance allele and drive-sensitive allele, and for moderate-effect drives and sufficiently low initial frequencies of the resistance allele, the drive allele essentially outran resistance to fixation. Once the drive fixed, type-M resistance had no effect.

The work here expands on the potential robustness of resistance-free evolution of unlinked suppression drives. Three meaningful elaborations can be offered:

- With homing drives, population suppression while avoiding resistance of type M can be increased by using multiple drives with individually mild effects instead of a single drive with the same total effect.
- Sib mating (type-P resistance) can evolve to suppress moderate-effect drives, but only if genotypes exist that enforce high levels of sib mating. High levels of sib mating can evolve despite being rare in the population, but low genetic levels of sib mating are not favored except with strong-effect drives. These results apply to both homing drives and toxin-antidote drives (here represented by ClvR).
- The realm of resistance-free evolution can be expanded by manipulating the ecological effects of the drive. Drives with density-dependent fitness effects can escape much of resistance evolution by externally imposed suppression of the population or by taking advantage of natural declines in population size (e.g., seasonality).

There are yet many elaborations to explore with the models, especially on the ecological end. Sib mating might increase dynamically (not through evolution) as the population size declines. Group selection itself (or its equivalent in a continuous population across space) may slow or even stop the spread of the drive (Bull et al. 2019; Champer et al. 2021). Our efforts here considered toxin-antidote drives (ClvR) in only a limited scope, and although there was nothing in our results to suggest a fundamental difference between resistance evolution for homing drives than for toxin-antidote drives, a more thorough consideration of toxin-antidote drives is warranted.

Despite the limited number of studies available, a distilled interpretation of them is that evasion of resistance evolution by a suppression drive is feasible but not assured. The most important consideration is to avoid ‘ allelic’ resistance, which for a homing drive would typically be a mutation in the nuclease cleavage site. There are now two feasible ways around that, one being to choose a cleavage site whose function is intolerant of mutation (Kyrou et al. 2018), the other being to take advantage of CRISPR to target multiple, independent sites in the same gene such that every chromosome in the population will have at least one site susceptible (Noble et al. 2017; Champer et al. 2018, 2020d; Edgington et al. 2020). The latter approach would greatly expand the opportunities for population suppression, but whether either approach will succeed may depend on successfully controlling expression of CRISPR proteins to avoid undesirable embryonic and somatic activity (e.g., Champer et al. 2017; Bier 2021; Hammond et al. 2021)). Other forms of type-M resistance, such as interference with nuclease expression and function, are not likely to be allelic or even closely linked to the drive, but chromosomal rearrangements could change linkage – to lock distant loci into a single linkage group. And if unlinked type-M and type-P resistance doesn’t already exist in the population, they are not likely to arise in time to stop a strong drive. However, deliberate introduction and expression of an anti-CRISPR protein, which may offer a fail-safe block against extreme suppression drives (Taxiarchi et al. 2021), may fail to block moderate-effect drives unless deliberately engineered to be closely linked to the drive or if introduced at high frequency.

Although suppression drives of moderate-effect may be the best bet for avoiding resistance evolution, they present some empirical difficulties. It is obvious that a suppression drive is only likely to succeed if it operates by destroying target gene function instead of carrying a deleterious cargo. Loss of a deleterious cargo would be a simple and evolutionarily unavoidable way to eliminate the suppressive effect of the drive. Furthermore, any genomic site chosen for disruption by a moderate-effect drive must be present in all members of the species, but it cannot be a strictly *essential* gene, for which null homozygotes would be inviable or sterile. This may raise the challenge of knowing the fitness effects of a null homozygote in advance of the release. Being assured of a fitness effect of 0.4 (for example) across the heterogeneous environment in the wild is not practical. Then, when the population persists after fixation of a moderate-effect drive (as will usually be the case), compensatory evolution can be expected to reduce the fitness effect of the knockout – type-C resistance Hall (2003); Rokyta et al. (2002); Harcombe et al. (2009); van Leeuwen et al. (2020).

Official releases of synthetic gene drives have yet to be approved. Any release of a suppression drive should anticipate the potential for resistance evolution at least because some types of resistance, once evolved, will thwart other attempts with suppression drives (e.g., type P). Thus the failure of one suppression drive may ensure the failure of subsequent drives. However, delaying the approval of official releases has its own risk in an age when multiple labs can create gene drives – the unintentional or unapproved deliberate release of a functional gene drive. There would seem to be some imperative for gaining experience with appropriately monitored gene drive releases before an uncontrolled release happens.

## Conclusion

A wide variety of gene drives can now be engineered, and experiments with model organisms in laboratory settings confirm their potential utility in suppressing populations. The unknown that faces releases into natural populations is the evolution of resistance. We previously showed that suppression drives with moderate fitness effects can evade many types of resistance that are unlinked to the drive locus. Here we have identified additional ways of escaping unlinked resistance. One method is to introduce multiple (e.g., two) drives of individually mild effect to achieve a greater combined suppression. Another method is to rely on the fact that resistance evolves in response to fitness effects of the drive; careful choice of ecological effects of the drive may thus enable population suppression with only modest fitness effects. One remaining challenge is that sib mating can evolve in response to a suppression drive with any fitness effect; sib mating can limit the suppression and even cause loss of the drive. However, sib mating is a threat for moderate-effect suppression drives only from alleles for high levels of sib mating (or if sib mating arises as a demographic effect of the reduction in population size). Evolution of resistance to suppression drives may therefore cause problems only on a limited basis.

## Data availability statement

The authors affirm that all information necessary for confirming the conclusions of the article are present within the article, figures, tables, and supplemental files.

The C programs used here are accessible at https://github.com/emuarmadillo/Gene_drives_C_Files_papers. The models calculate offspring genotype numbers by exhaustive enumeration of all possible parental matings multiplied by the fraction of gamete or offspring types produced by the parental genotypes; there are no explicit recursions of genotype frequencies in one generation from their frequencies in previous generations.

## Acknowledgments

We thank Jackson Champer for discussion and references. An anonymous reviewer prompted us to consider the effects of imperfect drive and heterozygote fitness effects and provided references. RG was supported by a WSU Honors College Distinguished Professorship, FC by a University of Idaho funds, and JJB by NIH grant GM 122079.

